# The landscape of DNA methylation associated with the transcriptomic network in laying hens and broilers generates insight into embryonic muscle development in chicken

**DOI:** 10.1101/470278

**Authors:** Zihao Liu, Xiaoxu Shen, Shunshun Han, Yan Wang, Qing Zhu, Can Cui, Haorong He, Jing Zhao, Yuqi Chen, Yao Zhang, Lin Ye, Zhichao Zhang, Diyan Li, Xiaoling Zhao, Huadong Yin

## Abstract

As DNA methylation is one of the key epigenetic mechanisms involved in embryonic development, elucidating its relationship with non-coding RNA and genes is essential for understanding early development. In this study, we performed single-base-resolution bisulfite sequencing together with RNA-seq to explore the genetic basis of embryonic muscle development in chicken. Comparison of methylome profiles between broilers and laying hens revealed that lower methylation in broilers might contribute to muscle development. Differential methylated region (DMR) analysis between two chicken lines showed that the majority of DMRs were hypo-DMRs for broilers. Differential methylated genes were significantly enriched in muscle development-related terms at E13 and E19. Furthermore, by constructing the network of the lncRNAs, we identified a lncRNA, which we named MYH1-AS, that potentially regulated muscle development. These findings reveal an integrative landscape of late period of embryonic myogenesis in chicken and give rise to a comprehensive understanding of epigenetic and transcriptional regulation, in skeletal muscle development. Our study provides a reliable data resource for further muscle studies.

## Introduction

Epigenetics mechanisms, including DNA methylation, histone modification, non-coding RNAs and chromatin remodeling, have been the subject of intense research over recent years because of their essential roles in various biological processes ^1,2^. These epigenetic mechanisms have been reported to be involved in human diseases^3^, oogenesis and spermatogenesis^4^ as well as in adipose and muscle development^5–7^. DNA methylation is an epigenetic mechanism that exerts considerable influence on the regulation of gene expression without changing the DNA sequence^8^. A role for DNA methylation in muscle development has been illustrated in human^9^, pig^5,6^, rabbit^10^, bovine^11^ and chicken^12^.

The embryonic stage is critical for muscle development in mammals, as the number of muscle fibers in the developing embryo remains stable after birth. Previous reports have demonstrated a function of DNA methylation in embryonic muscle development. For instance, Carrio et al.^13^ built the methylome of myogenic stem cells and demonstrated the importance of DNA methylation-mediated regulation of the cell-identity Myf5 super-enhancer during muscle-stem cell differentiation. Long noncoding RNAs have also been proven to be important in the regulation of muscle development. For example, linc-MD1 interacts with miR-133 and miR-135 to regulate the expression of transcription factors MAML1 and MEF2C that activate muscle-specific gene expression^7^. Recently, the regulatory relationship between DNA methylation and lncRNAs has drawn extensive research attentions and a database of methylation and lncRNA regulatory relationships has been built for human diseases studies^14^. However, studies on the role for this regulatory relationship in muscle development are limited. Zhang at el.^5^ reported the function of the lincRNA and DNA methylation regulatory relationship in muscle development in pig. Yang at el.^6^ revealed that DNA methylation potentially affects gene expression in skeletal muscle to influence the propensity for obesity and body size.

After long-term artificial breeding for different purposes, laying hens and broilers show great differences in the development of skeletal muscles. The skeletal muscle growth rate of broilers far exceeds that of laying hens even under optimal feeding conditions, and broilers can exhibit weights 5 times more than laying hens at 6 weeks of age. The comparatively similar genetic backgrounds and genomes of these two chicken lines allow for comparative studies of muscle development at the epigenetic level.

Several genome-wide methylation studies have been reported in chicken, and a relationship between DNA methylation level of promoters and expression level of genes were identified^15–17^. Furthermore, the global methylation landscape of muscle development was described in chicken using juvenile and later laying-period hens^12^. However, a role for DNA methylation in chicken embryonic muscle development has not been fully clarified.

Here we used whole genome bisulfite sequencing to determine the methylomes of 12 standardized broilers and 12 standardized laying hens. We sequenced the whole transcriptome of these 24 samples by RNA-seq simultaneously for the multi-Omics integrative analyses, to explore the effect of DNA methylation and lncRNA relationship on muscle development.

## Results

### Overview of DNA methylation

In the genomic methylation data among 24 samples (from 12 broilers and 12 laying hens), the average sequence depth is about 30.3X. Approximately 3.4 billion reads were generated by the Illumina HiSeq in total and an average of 71.99% clean reads were mapped to the *Gallus gallus* genome (version 5.0) (Supplementary Table S1). The coverage analysis revealed that approximately 82% of the *Gallus gallus* genome were covered by reads at least one-fold, whereas nearly 78% of genome was covered by more than five-fold and 75% of genome was covered more than 10-fold (Supplementary Table S2). These results indicated a reliable sequencing outcome.

The methylation level of each developmental stages is displayed in Fig 1a, which indicates that the layers and broilers have a similar global methylation profile. Similar proportions of CpGs in three sequence contexts (mCG, mCHG, and mCHH) were observed among four developmental stages (Fig. 1b). Next, the methylation level distributions of CpGs were analyzed at four developmental stages. In general, CpGs showed a high methylation level in the mCG context and a low methylation level in mCHG and mCHH contexts (Fig. 1c and Supplementary Fig. 1a). We then measured the methylation level of different regions of genes and compared these levels at different stages and populations. Interestingly, we found that broilers showed statistically lower methylation levels at all stages in the mCG context than layers (Fig. 1d). We quantified the numbers of CpG islands (CGIs) in different regions at different stages (Supplementary Fig. 1b). More CGIs were located in gene promoter regions in broilers than layers, which indicates that methylation in CGIs may be involved in faster muscle development in broilers, as CGIs located at promoter regions are important for controlling gene expression.^18^.

**Fig 1.**
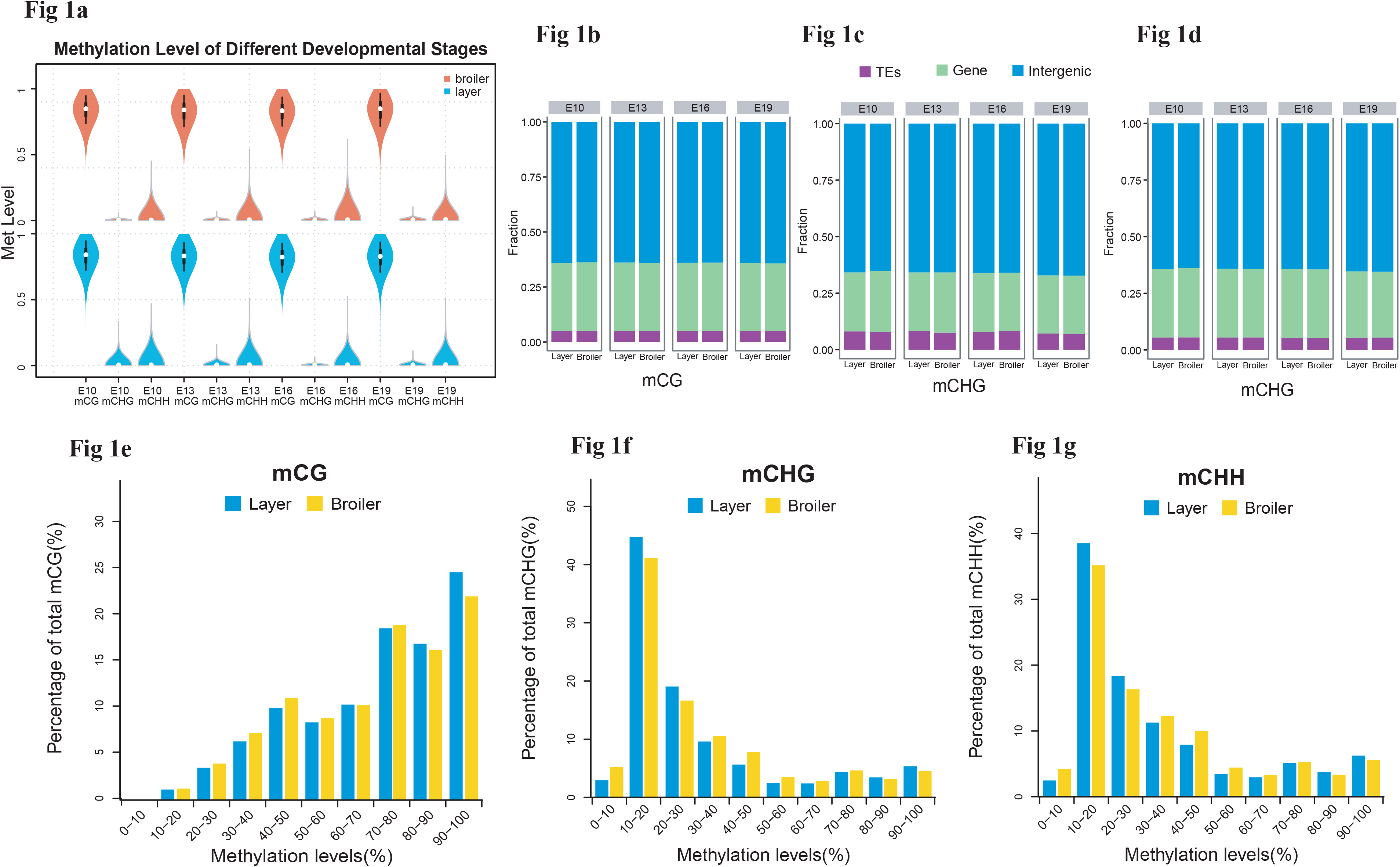
Genome-wide profiles of DNA methylation among different sample groups. (**a**) Genomic methylation level in either layers or broilers at E10, E13, E16, E19, respectively. Methylation level were range from 0 to 1. (**b-d**) Proportion of mCpG in different genomic features at different developmental stages in mCG, mCHG and mCHH contexts, respectively. (**e-g**) Methylation level of CpGs was equally divided into 10 intervals and the percentage of each interval were measured using E10 as example.

We also examined the methylation level of lncRNAs assembled in RNA-seq using a similar approach and compared levels with the analysis of gene methylations. Generally, broilers still showed a lower methylation level in various types of lncRNAs in mCG and mCHH contexts compared with layers; similar methylation levels were observed among different types of lncRNAs (Fig. 2b and Supplementary Fig. 2c-d). Genes and lncRNAs had similar global methylation levels and both showed significant difference in broilers compared with layers (Fig. 2a and Supplementary Fig. 2a-b). These results suggest that faster muscle development of broilers may be due to the lower methylation level in late embryonic stage compared with those in layers.

**Fig 2.**
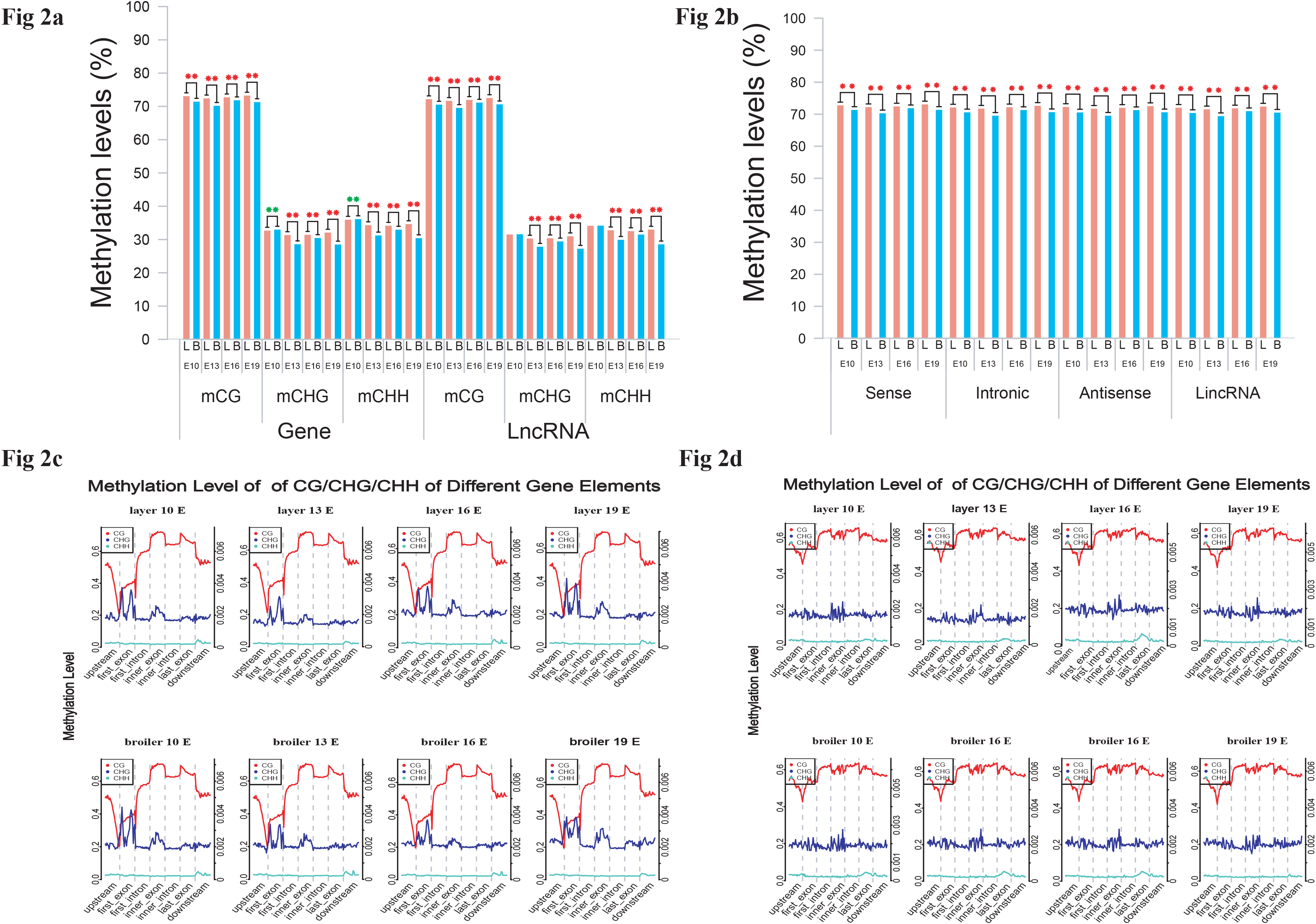
Comparatively measurement of methylation level of genes and lncRNA. (**a**) Comparison of methylation level of genes or lncRNAs between layers and broilers in three different contexts. (**b**) Measurement of methylation level of different types of lncRNAs. * *P* <0.05, ** *P* <0.01 for comparison between two chicken lines. The red star means the methylation level of layers is significantly higher than broilers whereas the green star represents an opposite result. (**c-d**) Genomic methylation around genes and lncRNAs were measured across the genome, respectively. Transcripts were separated into seven regions (upstream, first exon, first intron, inner exon, inner intron, last exon and downstream) and each region was equally divided into 20 bins for visualization.

We also analyzed the genomic distribution patterns of DNA methylation in genes and lncRNAs. We divided the upstream region (2 kb), first exon, first intron, internal exon, internal intron, last exon and downstream region (2 kb) of genes and lncRNAs across the genome as different features and their methylation levels were measured through 20 bins. In general, the 5' upstream and 3' downstream regions showed lower methylation levels than other gene regions. We also compared the methylation level of features of genes with features of lncRNA (Fig. 2c-d). LncRNAs have relatively higher methylation levels around the transcription start site (TSS) compared with genes (*P* < 0.001). In addition, methylation levels of different types of repeat regions were also analyzed across the genome. Beside the significant differences between broilers and layers, short interspersed nuclear elements (SINE) showed lower methylation levels across the four stages in the mCG context (Fig. 3 and Supplementary Fig. 3).

**Fig 3.**
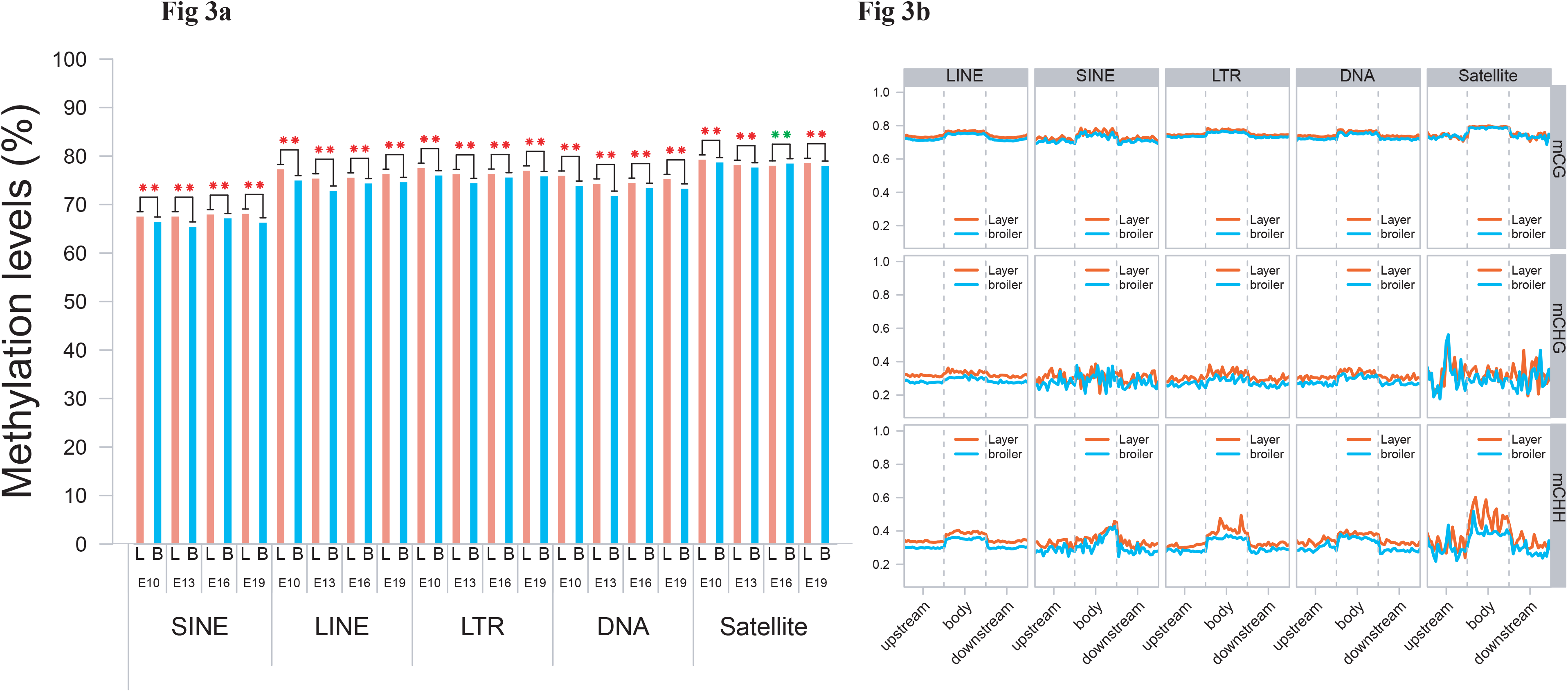
Methylation level of different types of TEs using E19 as an example. (**a**) Comparatively measurement of methylation of SINE, LINE, LTR, DNA, Satellite regions between two chicken lines in mCG context. (**b**) Methylation of different types of TEs for upstream, body and downstream regions in three different contexts using 20 bins across the whole genome.

### Identification of differential methylation regions (DMRs) and genes

To explore the potential causes of the divergence in muscle development between broilers and layers, the differential methylation loci were identified in DSS package. DMRs were identified in E10, E13, E16 and E19 based on differential methylation loci. The DMRs were subsequently annotated to the genome, and the distribution of the DMRs in the whole genome was analyzed (Fig. 4a and Supplementary Table S4-S7). In general, the majority of DMRs was located in intronic regions, and a small portion of DMRs was distributed in the promoters of genes (Fig. 4a). Proportion analysis revealed that broilers had more hypomethylated regions across the genome in the four developmental stages, indicating that low methylation in muscle development-related genes may account for the fast muscle development in broilers (Fig. 4b).

**Fig 4.**
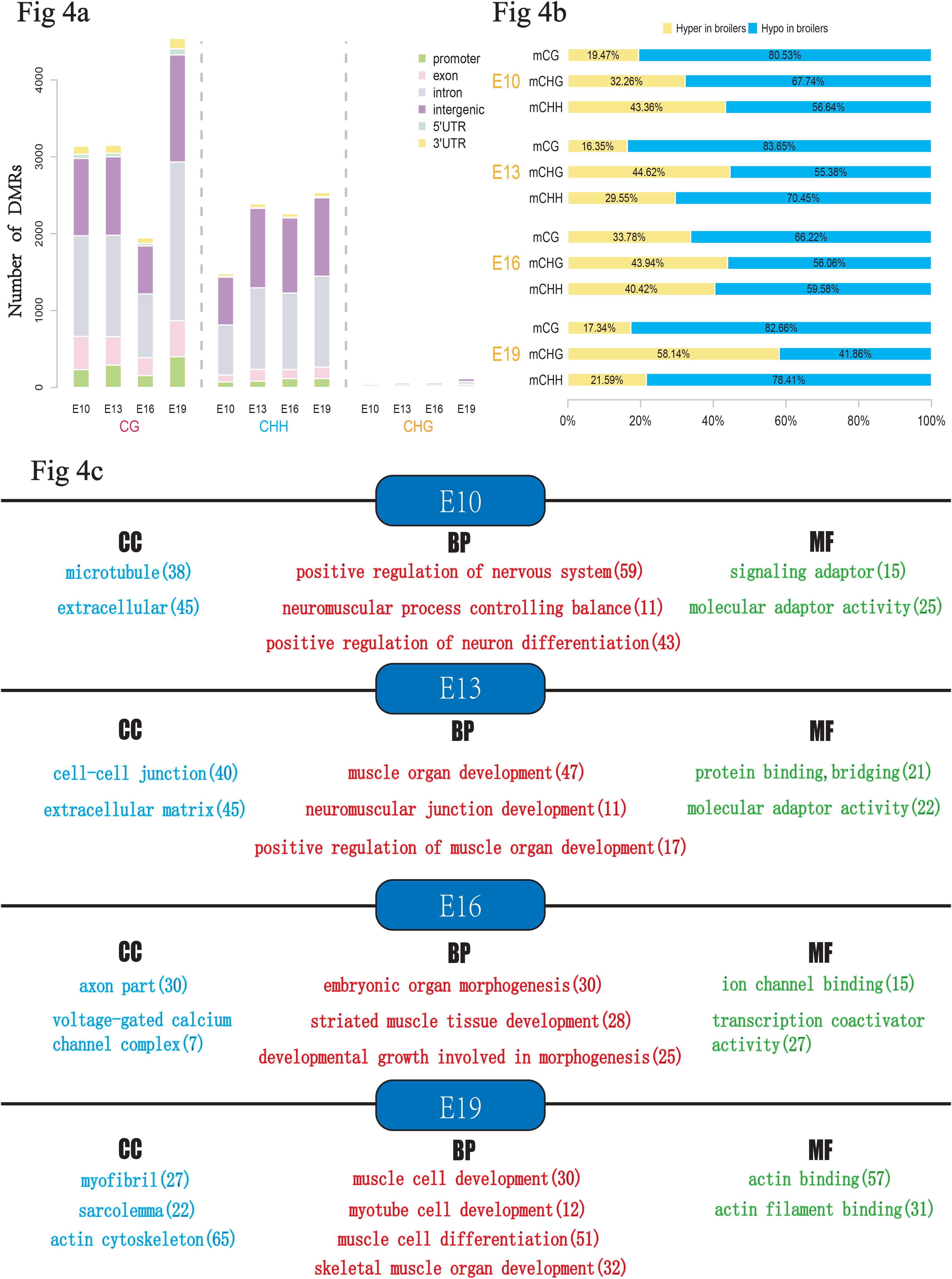
Analyses of DMRs at 4 developmental stages. DMR calling were performed in mCG, mCHG and mCHH, respectively. (**a**) Numbers of DMRs in different genomic features (promoter, exon, intron, intergenic, and UTR regions). (**b**) Relative proportion of hyper DMRs to hypo DMRs in different CpG contexts. (**c**) The results of Gene Ontology (GO) analysis for genes with overlapped with DMR. Only part of the terms was selected for display. The red color means GO-BP terms, the blue color means GO-CC terms whereas green color represents GO-MF terms. The number in bracket means number of genes enriched in a specific term.

Differential methylation genes (DMGs) were defined as genes with at least one overlapping DMR in its exon/intron regions. Gene Ontology (GO) enrichment analyses were then performed to investigate potential biological functions of the DMGs. In general the DMGs in the four developmental stages were most significantly enriched in terms related to the nervous system. However, many muscle-related terms were also found, especially for DMGs at E13 and E19, such as muscle organ development (47 genes; Q-value < 0.001), myotube cell development (12 genes; Q-value < 0.005), positive regulation of muscle organ development (17 genes; Q-value < 0.001), and muscle cell differentiation (51 genes; Q-value < 0.003) (Fig. 4c, Supplementary Table S8-S11). Because DMRs were not unanimous on the genomic position among different developmental stages, we merged the genomic position of DMRs from the 24 samples to generate common DMRs and re-calculated the methylation level for each common DMR. Clustering analysis was performed using the common DMRs and displayed using heatmap analysis. Different developmental stages were shown to cluster together, which is indicative of the high quality of sampling and DMR calling in this experiment (Fig. 5a). Moreover, the principle component analysis (PCA) result was consistent with the clustering analysis (Fig. 5b).

**Fig 5.**
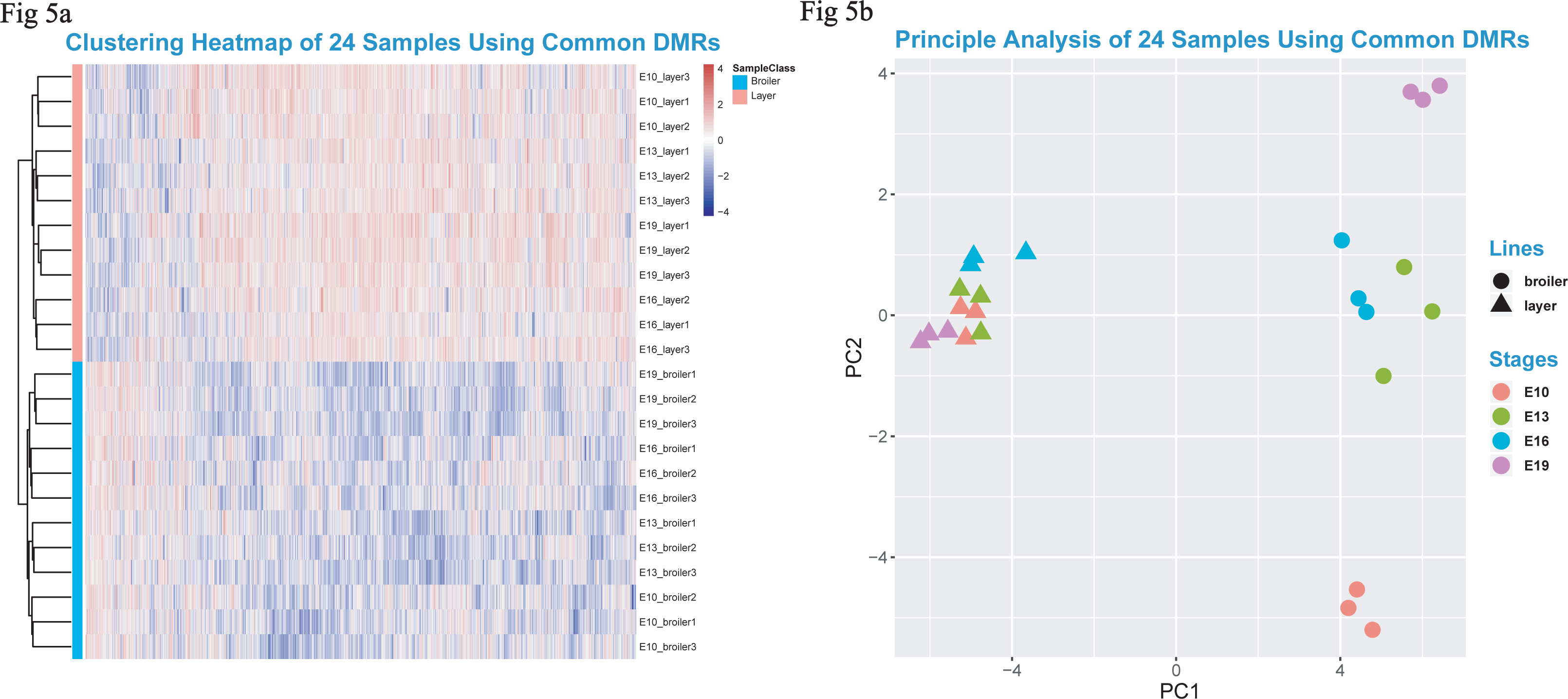
Heatmap clustering analysis and PCA analysis. (**a**) Heatmap clustering using merged common DMRs among 24 samples (see Materials and Methods). (**b**) The result of PCA analysis using common DMRs among 24 samples. Only the first component and the second component were visualized.

### Integrative analyses of DNA methylation and transcriptome

To further explore whether methylation influences gene and lncRNA expression in chicken, RNA-seq was used to measure the expression of genes and identified lncRNAs. We identified 20656 lncRNAs in total. Most of the lncRNAs were lincRNAs (63.6%) (Fig. 6a, 6b). Heatmap of 24 samples and PCA suggested developmental stages accounted for most variances (Fig. 6c). We divided genes and lncRNAs into four groups on the basis of their expression level (highest, medium high, medium low and lowest) using quantile method. We then measured methylation levels in different groups of genes and lncRNAs. In general, broilers and layering hens had similar methylation levels. A negative correlation was observed between genes and methylation of promoters in both broilers and layers: the highest expression level group showed the lowest methylation level around the TSS, whereas the lowest expression level group showed the highest methylation level (Fig. 6d, e). Interestingly, the trend of negative correlation between expression and methylation was observed in downstream regions of lncRNAs but not around the TSS (Fig. 6f, g). Moreover, the lncRNAs were usually methylated at higher levels around the TSS compared with genes (Fig. 6d-g).

**Fig 6.**
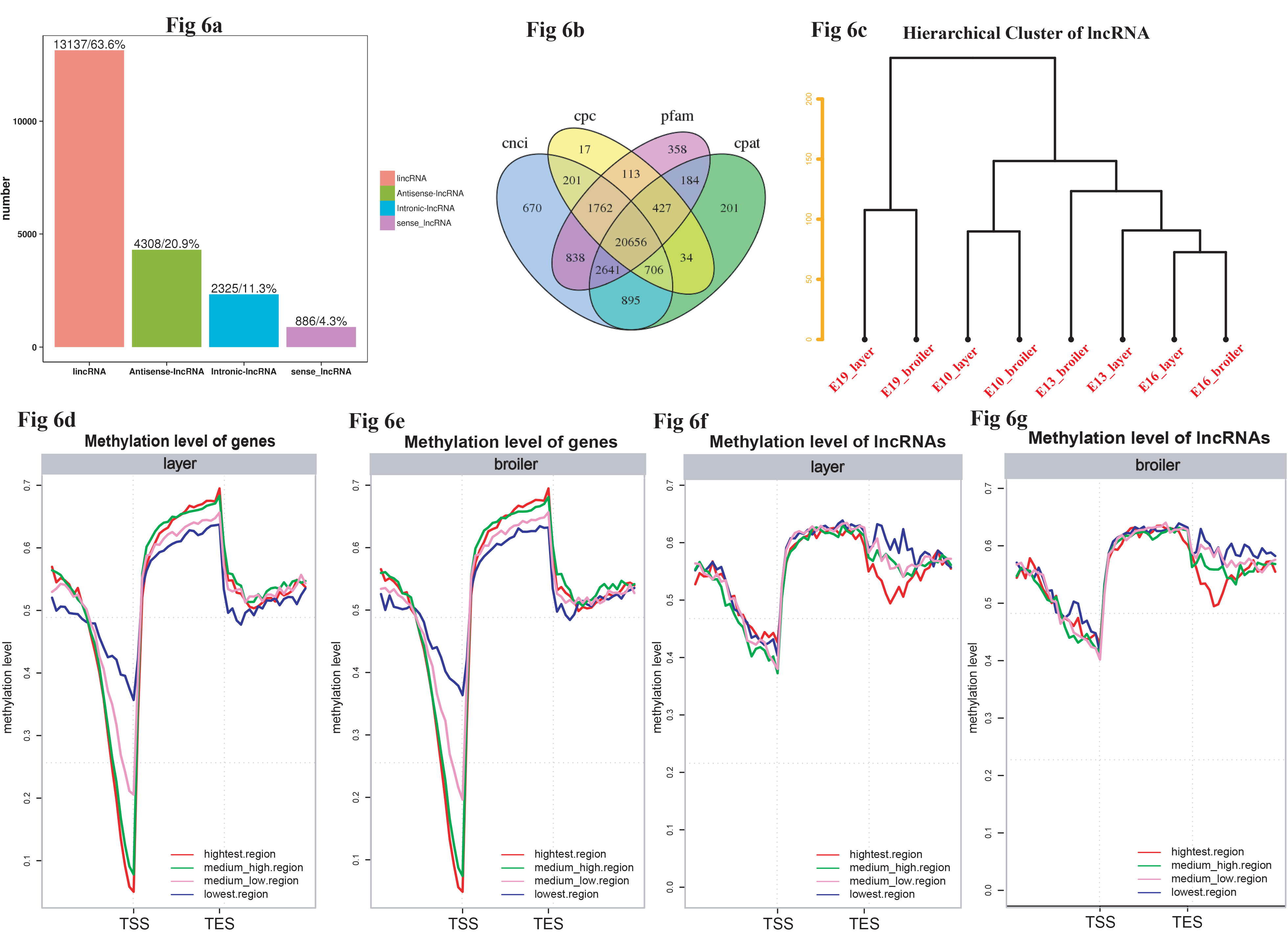
LncRNAs idenditication and correlation analysis between methylome and transcriptome. (**a**) Number of different types of lncRNAs in all developmental stages. (**b**) Venn diagram of lncRNAs identified through different software. (**c**) Hierarchical cluster analysis of lncRNAs using their expression level. Replicates were merged together in the analysis. (**d-g**) The genes and lncRNAs were divided into five groups based on their expression levels, respectively. Then the methylation level around TSS and TES of each group were measured using 20 bins across the whole genome for layers and broilers.

Next, differential expression gene (DEG) and lncRNA (DEL) calling was performed, and the cis-targets and trans-targets of lncRNAs were predicted. The DMRs were assigned to lncRNAs generated from RNA-seq in this study (Supplementary Table S12-S15) and the differential methylation lncRNA (DM lncRNA) were defined as DEL that overlapped with DMR. The result showed that 55 DM lncRNAs were identified (13,16,11,15 in 4 stages, respectively) (Supplementary Table S16). We then searched for DM lncRNAs with potential in regulating muscle development. In particular, we found that the expression of one lncRNA (which we named as MYH1-AS; Fig. 7a) was highly correlated with the methylation level of the DMR assigned to it (Spearman, Cor=-0.7513, *P* < 10^-4^; Fig. 7b). The expression of MYH1-AS was detected to dramatically increase in broilers compared to laying hens at E16 and E19 (Fig 8a). As the lncRNA was predicted by by lncTar to target several genes like MYH1A, MYH1G and MYH1E, the expression correlations between the lncRNA and its targets were calculated to search for its most likely target. MYH1E showed the highest correlation with MYH1-AS (Fig. 7d), indicating MYH1E as a potential target of MYH1-AS. To further explore the role of MYH1-AS in muscle development, the gene-lncRNA networks were constructed based on their mRNA expression connectivity using WGCNA, and the subnetwork of MYH1-AS was extracted from the whole network. MYH1-AS had a high correlation with several muscle-related genes in this subnetwork (Fig. 7d). The relationship between the connectivity and correlation is shown in Figure 7f. Interestingly, genes that were highly negatively correlated with MYH1-AS did not show high connectivity with MYH1-AS. All genes showing high connectivity with MYH1-AS were also highly positively correlated with the lncRNA (Fig 7e-f). A total of 168 genes with both high connectivity and correlation with MYH1-AS, were selected to perform GO enrichment analysis to confirm the role of MYH1-AS in muscle (Fig. 7g and Supplementary S17). The results showed that the majority of terms enriched by these genes were muscle-related.

**Fig 7.**
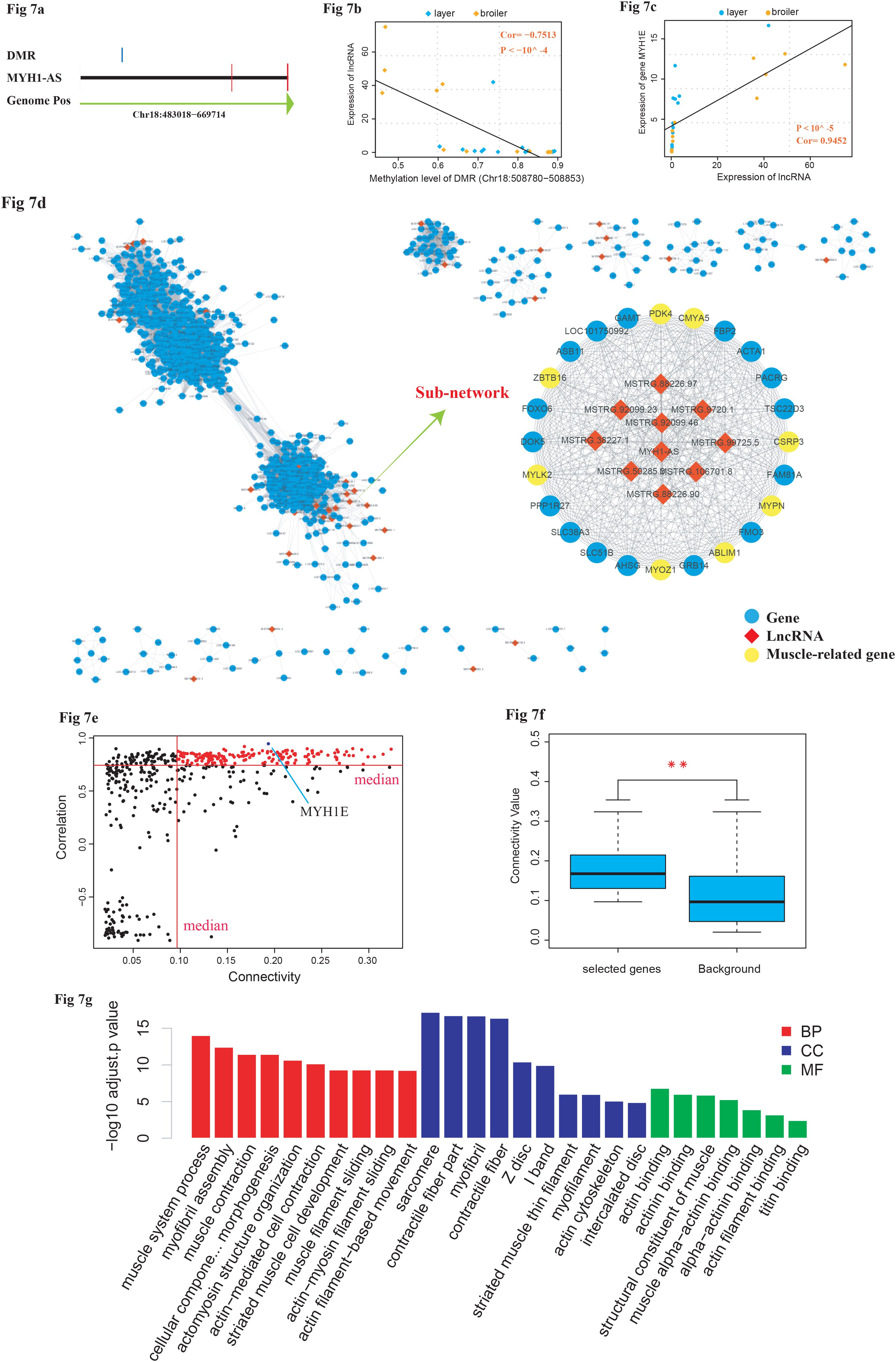
Comprehensive analysis of lncRNA MYH1-AS. (**a**) visualization of the transcript of MYH1-AS and DMR overlapped it. (**b**) Correlation between methylation of DMR and expression of MYHA-AS using Spearman method. (**c**) Correlation between expression of MYH1-AS and expression of its potential target MYH1E. (**d**) The whole gene-lncRNA network and subnetwork including MYH1-AS extracted from the entire network. (**e**) Relationship between correlation and connectivity of gene and MYH1-AS. The red points represent genes with both high connectivity and correlation with MYH1-A and were selected for subsequent GO analysis. (**f**) Comparison of connectivity value between genes selected (red points) and all genes with in the subnetwork (background). * *P* <0.05, ** *P* <0.01 for comparison between selected genes and background. (**g**) Results of GO analysis for genes selected.

The expressions of MYH1-AS produced by RNA-seq were verified by qPCR and a similar trend was observed, indicating a reliable sequencing outcome (Fig 8 a, b). Subsequently, a siRNA was designed to perform MYH1-AS silencing assay. As shown in fig 8c, expression of MYH1-AS was significantly reduced after transfecting, indicative of efficiency of siRNA used in this experiment (Fig 8c). Then the mRNA expression of muscle related genes (MyoD1, MyoG and MyH3) were measured at 48h after MYH1-AS silencing. It resulted in a reduced mRNA expression in silencing groups compared to control groups (Fig 8d-f). Besides, the microscope was used to monitor the morphological change in myotubes after silencing. We found that MYH1-AS silencing resulted in a reduced number of myotube (Fig 8g-h). Further western blot assay revealed that the protein expression of MyhC and MyoG was repressed in silencing groups (Fig 8i). Those results suggest that lncRNA MYH1-AS may function in muscle differentiation.

**Fig 8.**
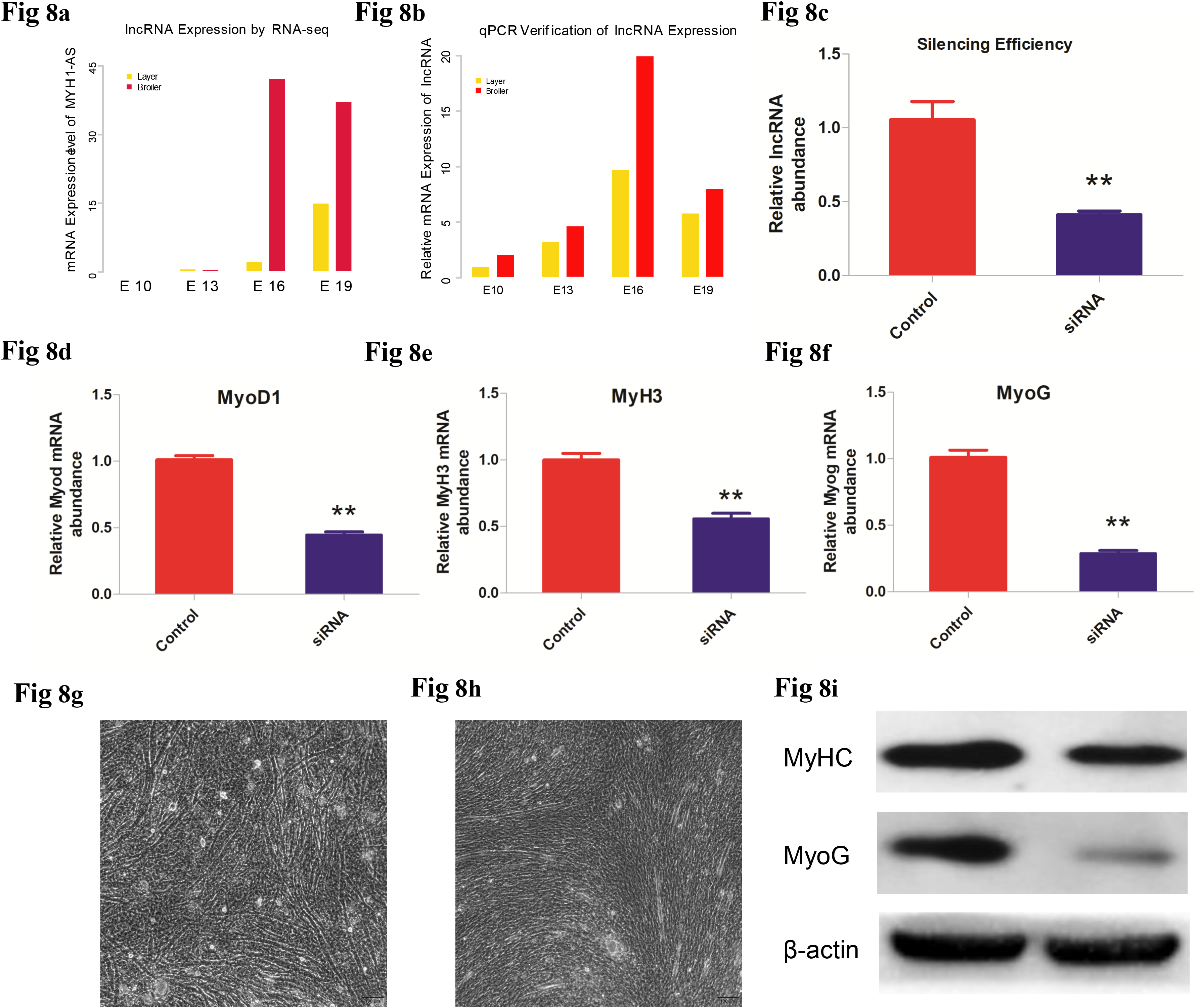
(**a**) Expression level of MYH1-AS in layers and broilers at different developmental stages. (**b**) Verification of lncRNA MYH1-AS expression at four developmental stages by qPCR. (**c**) lncRNA Silencing efficiency. * *P* <0.05, ** *P* <0.01 for comparison between control and silenced group. (**d-f**) The mRNA expression of MyoD1, MyH3 and MyoG in control and MYH1-AS silenced groups, respectively. * *P* <0.05, ** *P* <0.01 for comparison between control and silenced group. (**g-h**) The morphological changes in myotubes after silencing. (**8i**) The protein expression of MyHC and MyoG comparison between control and silenced group, respectively.

## Discussion

The chicken provides a unique model to perform embryology research because of the accessibility of egg. As chicken is an important food source for the human diet, the muscle development of chicken is an important topic worth of study. Here we used broilers and laying hens to explore the muscle development in chicken in the late embryonic period as they are artificially selected for different commercial use (depositing meat and laying eggs, respectively) thereby are divergent in muscle development. Because of the crucial role of methylation in embryogenesis, we performed whole genome bisulfite sequencing and RNA-seq to systematically explore the prenatal methylation landscape during chicken muscle development. Previous methylome studies have been performed using prenatal chicken or born chicken muscle^12,19,20^, however, these studies failed to generate a comprehensive methylation landscape of embryonic stages. We focused on more systematical study at embryonic stage range from E10 to E19 between two chicken lines and aimed to elucidate the detain of embryonic muscle development.

The methylation level and proportion of different methylations (mCG, mCHG, mCHH) of each developmental stage indicated that layers and broilers have a similar global methylation profile. We also measured the methylation level of different types of CpG (Fig. 1e-g), and results were consistent with previous studies in chicken muscle^15^. The distribution proportions of CpG in the genome were different from those in the study of Zhang et al^20^, as the CpG proportions in repeat regions accounted for less genomic proportion in our study. One possibility for the discrepancy may be because the previous study used data from born chicken, whereas our analyses were performed in data from prenatal chicken. More studies are required to clarify these differences.

We next comprehensively compared the methylation level of genes and lncRNAs among different developmental stages and chicken lines (Fig. 2a). In general, layers showed a significantly higher methylation level than broilers in the mCG context in both genes and lncRNAs, which may be responsible for the differences in muscle development. Furthermore, we compared the methylation levels of different types of lncRNAs (sense, intronic, antisense and lincRNA) and there were no significant differences, although layers and broilers still revealed significant variances. Next, genomic methylation around genes and lncRNAs were measured across the genome, and the TSSs were found to be low methylated in genes (Fig. 2c). The broilers and layers showed similar trends around the TSS, which is consistent with patterns reported in previous studies in chicken^12,15^, as well as in bovine muscle tissue^11^ and pig^21^. However, the TSSs of lncRNAs were usually methylated at higher levels compared with genes, which may explain why mRNA expression of lncRNAs are usually lower than genes (*P* < 10^-8^) because methylation events in the promoter region usually affect gene expression^22^. In addition, the methylation levels of different types of transpose elements (TEs) (SINE, LINE, LTR, DNA and satellites) were also measured and TEs were methylated at higher levels in layers compared with broilers. TEs are usually inactivated in animals but were reported to function in the early development of human and other mammals to provide cis-regulatory elements that coordinate the expression of groups of genes^23^. As epigenetic regulation is important for the activity of TEs^24^, these differences in the two chicken lines may also account for the divergence in development.

The clustering heatmap and PCA were performed using common DMRs among four developmental stages. The expected classifications were observed in both analyses, indicating the reliable outcomes of sequencing and DMR calling. Moreover, we found that DMRs between two chicken lines mainly distributed in intron regions and intergenic regions. These results are consistent with previous studies in chicken^12^, indicative of the important role of methylation in development regulation. However, as methylation in gene body region affects gene expression in several sophisticated ways^18^, further studies on how methylation of the intron regions can influence gene expression are required to elucidate the complicated epigenetic mechanism underlying muscle development in chickens. We analyzed the proportion of hypermethylated and hypomethylated regions and the majority of DMRs were detected to be hypomethylated regions in broilers, indicating that low methylation may be responsible for fast muscle development. This result is consistent with the preceding results in this study. Genes with overlapped with DMR at different times were regarded as DMGs and used for GO enrichment analysis. We found that DMGs at E13 and E19 were significantly enriched in muscle-related terms, suggesting that methylation plays an important role in embryonic stage muscle development. Additionally, DMGs among four stages were significantly enriched in nerve development-related terms, which may relate to the impact of domestication and artificial breeding. Integrative analysis was conducted to study the association between methylation level and mRNA expression. We noticed that mRNA level and methylation level around TSSs were negatively correlated in genes but not lncRNAs, indicating that DNA methylation regulates lncRNA expression in a more complex way than gene expression.

To explore which lncRNA may potentially influence muscle development, the DM lncRNAs were identified and the correlation between DM lncRNA and the assigned DMR were measured. In particular, MYH1-AS showed a high correlation with its target MYH1E and the DMR located in its intron region. Further WGCNA analysis revealed that several muscle-related genes were highly correlated with MYH1-AS in its subnetwork. For example, *MYLK2*, a muscle-specific gene, expresses skMLCK specifically in skeletal muscles^25,26^. ABLIM1 was reported to be related to muscle weakness and atrophy^27^. Increased PDK4 expression may be required for the stable modification of the regulatory characteristics of PDK observed in slow-twitch muscle in response to high-fat feeding^28^, and other genes in the network, such as MyoZ1, MYPN and ZBTB16 genes, were also revealed to be muscle- or meat quality-related genes^29–32^. This indicates that MYH1-AS may function in muscle development. Notably, as we noticed that high correlation did not exactly indicate high connectivity (Fig. 7f), we also performed GO enrichment analysis using 168 genes, which had top 50% both high connectivity and correlation values with MYH1-AS in its network as input. The majority of the resulting GO terms were muscle-related terms (Fig. 7f-g), which is strongly indicative of MYH1-AS functioning in muscle development. Therefore, these results suggest that MYH1-AS is regulated by DNA methylation and participates in muscle development during embryonic stages. Subsequent silencing and western blot assay verified our analysis results, suggesting the reliability of our analysis and the role of MYH1-AS in muscle differentiation. However, how the lncRNA regulates muscle development requires more studies.

Our experiment revealed a comprehensive DNA methylome and transcriptome landscape during embryonic developmental stages. We identified one lncRNA, MYH1-AS, that may potentially play a part in muscle development in chicken, and our study provides evidence for this conclusion. Moreover, we provided a basis and a reliable resource for further investigating the genetic regulation of methylation and gene expression in embryonic chicken. However, more studies are needed to elucidate the detailed mechanism on how DNA methylation impacts lncRNA expression and how the lncRNA regulates myogenesis.

## Acknowledgements

We thank Edanz Group (www.edanzediting.com/ac) for editing a draft of this manuscript.

## Materials and Methods

### Sample collection

The fertilized eggs of Rose and WhiteLoghorn were incubated in the same condition. The breast muscle and blood were collected at E10, E13, E16, E19. After sex determination, only samples identified as male were kept for next experiment. A total of 24 embryonic chicken were used in the study to form eight groups: E10, E13, E16, E19 for Rose and WhiteLoghorn, respectively. Each group included 3 individuals as biological replicates.

### DNA and RNA extraction

Genomic DNA was extracted using an animal genomic DNA kit (Tiangen, China) following the manufacturer’s instructions. The DNA integrity and concentration were measured by agarose gel electrophoresis and NanoDrop spectrophotometer, respectively. Total RNA was isolated using TRIzol (TAKARA, Dalian, China) 110 reagent according to the manufacturers’ instruction. RNA was reverse 111 transcribed by TAKARA PrimeScriptTM RT reagent kit (TAKARA) 112 according to the manufacturers’ instruction.

### Library construction and sequencing

Bisulfite sequencing libraries were prepared using the TruSeq Nano DNA LT kit (Illumina, San Diego, CA, USA). The genomic DNAs were then fragmented into 100-300 bp by sonication (Covaris, USA) and purified using a MiniElute PCR Purification Kit (QIAGEN, Silicon Valley Redwood City, CA, USA). The fragmented DNAs were end repaired and a single 'A' nucleotide was appended to the 3' end of each fragment. After ligating the DNAs to the sequencing adapters, the genomic fragments were bisulfite converted via a Methylation-Gold kit (ZYMO, Murphy Ave. Irvine, CA, USA). The converted DNA fragments were PCR amplified and sequenced as paired-end reads using the Illunima HiSeq xten platform by the Biomarker Technologies company (Beijing, China).

### Data alignment and process

The raw data in the FastQ format generated by the Illumina HiSeq were pre-processed by removing reads containing adapters, N (unknown bases) > 10%, and those which over 50% of the sequence exhibited low quality value (Qphred score ≤ 10). During the process, we also calculated the Q20, Q30, CG content for each sample data. The reads remained after this procedure were clean reads and used for subsequent analysis. The methylation data were aligned to reference genome Gallus gallus 5.0 by Bismark software^33^. Meanwhile, the number of aligned clean reads in unique position of reference genome were calculated as unique mapped reads number. The proportion of the number of aligned reads in the total number of reads was calculated as the mapping rate. Subsequently, the methylation level of single base was then calculated by the ratio of the number of methylated reads to the sum of total reads covered the locus. Finally, we used a binominal distribution teat approach to determine whether a locus was regarded as methylated locus with the criteria: coverage depth > 4 and FDR<0.05^33^.

The transcriptional libraries were sequenced on an Illumina HiSeq xten platform at the Biomarker Technologies Company (Beijing, China). The obtained transcriptome data were filtered by removing sequences containing adaptors, low-quality reads (Q-value < 20), and reads containing more than 10% of unknown nucleotides (N) and were aligned to reference genome Gallus gallus 5.0 by HISAT2^34^ then the transcript assembly and FPKM calculation were performed using the StringTie^35^. Transcripts mapped to the coding genes of reference were used to subsequent differential expression gene calling.

### LncRNA identification

In order to identify the potential lncRNA, the assembled transcripts generated from the StringTie were submitted to CPC^36^, CNCI^37^, CPAT^38^ and pfam^39^ software with defeat parameters to predict the potential lncRNAs. Only transcripts predicted as lncRNA shared among four tools were regarded as candidate lncRNA. Then the cis-target gene of lncRNA were defined as neighbor gene in 100 kb genomic distance from the lncRNA and were identified using in-house script. The trans-target prediction of lncRNAs was performed by LncTar software^40^.

### DMLs and DMRs calling

The differential methylation locus (DMLs) and differential methylation regions (DMRs) between broilers and layers at each comparison were detected separately using Dispersion Shrinkage for Sequencing Data (DSS) package in R^41–44^. The differential methylation regions (DMRs) were then calculated in with default parameters. Subsequently, DMRs were annotated using ChIPseeker package in R^45^.

Gene overlapped with at least one DMR is defined as differential methylation gene (DMG). Common DMRs among 4 developmental stages were identified by merging all positions of DMRs in 24 samples and re-calculating the methylation level for each merged DMR position with an average approach using mCpG data.

### DEGs and DELs calling

The differential expression genes (DEGs) calling and the differential expression lncRNA (DEL) calling between two populations at each time point were performed separately using the DEseq^46^. The results were filtering with the criteria: (1) fold change >2 (2) FDR<0.5. The transcripts satisfied both standards were regarded as DEGs or DELs.

### Functional enrichment analysis and WGCNA analysis

Gene ontology enrichment analyses were conducted for DMGs at E10, E13, E16, E19 comparisons respectively to explore their potential roles in muscle development. These analyses were performed by clusterProfiler package implemented in R^47^. A hypergeometric test was applied to map DMGs to terms in the GO database to search for significantly enriched terms in DMGs compared to the genome background.

The WGCNA analysis was performed using WGCNA package implemented in R <u>ENREF 48</u>^48^. We used all the differential expression lncRNAs and all the genes as input. Then, variable coefficient was used to filter transcripts with low expression change. The variable coefficient was calculated as follow: C_v_ =σ/μ. The σ is the standard deviation and μ represents the mean value of expression of input transcripts. Only transcripts with ranked top 30% high C_v_ value were used for WGCNA analysis. After the entire network was constructed, only genes with connectivity more than 0.15 were selected for subsequent subnetwork analysis.

### Validation for RNA-seq by quantitative Real-time RCP(Q-PCR)

Total RNA was purified and reversely transcribed into cDNA using PrimerScriptR RT reagent Kit with gDNA Eraser (Takara Biotechnology (Dalian) Co., Ltd) following the specification. Quantities of mRNA were then measured with qRT-PCR using a CFX96TM real-time PCR detection system (Bio-Rad, USA). The qRT-PCR assays were then performed with a volume of 20 μL containing 10 μL SYBR Green Mixture, 7 μL deionized water, 1 μL template of cDNA, 1 μL of each primer and with following thermal conditions: 95 °C for 5 min, 45 cycles of 95 °C for 10 sec, 60 °C for 10 sec, 72 °C for 10 sec. Primer sequences used for qRT-PCR assays are displayed in Supplementary Table 17. β-actin gene was used as internal control. Each qPCR assay was carried out in triplicate. The relative gene expression was calculated by using the 2-ΔΔCt method.

### Cell cultures

Post-hatch chickens (7-day-old commercial generation Avian broiler chicks) were purchased from Wenjiang Charoen Pokphand Livestock & Poultry Co., Ltd. The pectoralis muscle was removed and used for preparation of primary myogenic cultures. About 5 g of muscle was finely minced and treated with 0.1% collagenase I (Sigma, MO, USA) followed by 0.25% trypsin (Hyclone, UT, USA) to release cells. Then, the cell suspension was subjected to Percoll density centrifugation to separate myoblasts from contaminating myofibril debris and nonmyogenic cells. Cells were plated in 25 cm3 cell culture bottles with complete medium [DMEM/F12 (Invitrogen, Carlsbad, CA) +15% FBS (Gibco, NY, USA) +10% horse serum (Hyclone, UT, USA) +1% penicillin-streptomycin (Solarbio, Beijing, China) +3% chicken embryo extraction]. The cells cultured at 37 °C and 5% CO2 with saturating humidity, which were allowed to proliferate in growth medium for 2-4 d, and the medium was refresh every 24 h. To induce differentiation, satellite cells were grown to 80% confluence in growth medium, and the replaced with differentiation medium composed of DMEM, 2% horse serum and 1% penicillin-streptomycin, and the medium was refreshed every 24 h.

### LncRNA silencing

Chicken satellite cells were cultivated in 6-well plates and transfected with siRNAs: 5'-GGAAGGGAGUAGGUGGUAATT-3' and 5'-UUACCACCUACUCCCUUCCTT-3'; Sangon Biotech, Shanghai, China) when grown to a density of approximate 70% in plates. In contrast, control cells were transfected with negative siRNA with same other condition. The transfection reagent was Lipofectamine 3000 (Invitrogen, Carlsbad, CA, USA). The knockdown efficiency was assessed by quantitative RT-PCR of lncRNA MYH1-AS.

### Microscopy

Cellular morphology was evaluated in differentiated myotubes by phase-contrast microscopy without preliminary fixation. Pictures were produced using the Olympus IX73 inverted microscope (OLYMPUS, Tokyo, Japan) and the Hamamatsu C11440 digital camera (HAMAMATSU, Shizuoka, Japan).

### Western blot assay

The cells were collected from the cultures, placed in the RIPA lysis buffer on ice (BestBio, Shanghai, China). The whole proteins were subjected to 10% sodium dodecyl sulfate polyacrylamide gel electrophoresis (SDS-PAGE) and then transferred to polyvinylidene fluoride membranes (PVDF; Millipore Corporation, Billerica, MA, USA). The PVDF membrane was incubated with 5% defatted milk powder at room temperature for 1 h, then incubation with the following specific primary antibodies at 4°C overnight: anti-MyoG (Abcam), anti-MyHC (Abcam) and anti-β-Actin (Abcam). The secondary antibodies HRP-labeled rabbit IgG (Cell Signaling) were added at room temperature for 1h. Following each step, the membranes were washed five times with PBS-T for 3 min. The proteins were visualized by enhanced chemiluminescence (Amersham Pharmacia Biotech, Piscataway, NJ, USA) with a Kodak imager (Eastman Kodak, Rochester, NY, USA). Quantification of protein blots was performed using the Quantity One 1-D software (version 4.4.0) (Bio-Rad, Hercules, CA, USA) on images acquired from an EU-88 image scanner (GE Healthcare, King of Prussia, PA, USA).

